# Non-destructive, whole-plant phenotyping reveals dynamic changes in water use efficiency, photosynthesis, and rhizosphere acidification of sorghum accessions under osmotic stress

**DOI:** 10.1101/2023.09.26.559576

**Authors:** Daniel N. Ginzburg, Jack A. Cox, Seung Y. Rhee

## Abstract

Noninvasive phenotyping can quantify dynamic plant growth processes at higher temporal resolution than destructive phenotyping and can reveal phenomena that would be missed by end-point analysis alone. Additionally, whole-plant phenotyping can identify growth conditions that are optimal for both above- and below-ground tissues. However, noninvasive, whole-plant phenotyping approaches available today are generally expensive, complex, and non-modular. We developed a low-cost and versatile approach to non-invasively measure whole-plant physiology over time by growing plants in isolated hydroponic chambers. We demonstrate the versatility of our approach by measuring whole-plant biomass accumulation, water use, and water use efficiency every two days on unstressed and osmotically-stressed sorghum accessions. We identified relationships between root zone acidification and photosynthetic efficiency on whole-plant water use efficiency over time. Our system can be implemented using cheap, basic components, requires no specific technical expertise, and is suitable for any non-aquatic vascular plant species.

## Introduction

Water and nutrient availability, along with the efficiency of their use by crops, are major determinants of agricultural productivity (Fageria et al., 2007; Sinclair and Rufty, 2012). However, unchecked demand for natural resources, such as land, fertilizer, and water, to support global agricultural production has led to widespread ecosystem degradation and biodiversity loss (Foley et al., 2005; Williams et al., 2020). Thus, the demand for stable agricultural yields amidst globally changing climate and decreasing natural resource availability necessitates more resource-efficient and climate-resilient crops and agronomic practices.

Plant growth is neither linear over time (Paine et al., 2011) nor uniform across genotypes within a species (Mortlock and Hammer, 1999; Sugiyama et al., 2012). Measurements taken at single time points thus fail to capture the dynamic variation of plant growth across developmental stages and environmental conditions (Granier and Vile, 2014). In addition to being generally higher throughput than destructive measurements, noninvasive phenotyping approaches provide a window into temporal dynamics along a developmental trajectory and in response to changing environmental conditions. Immense progress has been made in high-throughput phenotyping to capture plant traits non-invasively, particularly through the use of advanced imaging platforms (Shakoor et al., 2017; Li et al., 2020). For example, by non-invasively tracking growth over time in various *Setaria* spp., in which visual leaf area strongly correlates with above-ground biomass, Fahlgren and colleagues uncovered species-specific growth dynamics across time and environmental conditions which would have otherwise been missed by endpoint measurements alone (Fahlgren et al., 2015).

In contrast to phenotyping approaches that capture data from only certain plant parts, whole-plant phenotyping can identify phenotypes and growth conditions that are optimal for both above- and below-ground tissues, and is thus likely more valuable for breeding more resilient and resource-efficient crops (Chochois et al., 2015). Because of the relative difficulty and time required to measure traits at the whole-plant level (Li et al., 2020), measurements are often simplified to include only above-ground plant tissue. However, approaches that exclude root biomass are potentially limited in value. For example, identifying genetic variation in traits such as biomass accumulation and water use efficiency is particularly important in the context of water-limited conditions in which the ratio of root:shoot biomass is often substantially different than under water-replete conditions (Hubick et al., 1986; Benjamin et al., 2014; Xu et al, 2015). Even in well-watered conditions, the ratio of root:shoot biomass partitioning can be different among genotypes (Chochois et al., 2015; Chenu et al., 2018), further highlighting the importance of including below-ground biomass to accurately quantify resource use efficiency (Leakey et al., 2019).

Various approaches can phenotype below-ground traits non-invasively. For example, root growth can be measured *in vivo* using magnetic resonance imaging (MRI) (Rascher et al., 2011), X-ray computed tomography (CT) scanning (Zeng et al., 2021) or with luminescence-based reporters (Rellán-Álvarez et al., 2015). When combined with non-destructive techniques to measure above-ground tissues, such approaches could in theory measure whole-plant growth over time. However, these approaches often require large upfront costs, state-of-the-art technologies, or specific technical expertise (Großkinsky et al., 2015). There are reports of cheaper and simpler approaches to measure whole-plant traits non-destructively (Fletcher et al., 2018). However, the authors found that data obtained non-destructively did not correlate well with values from plants harvested for end-point measurements.

To address the various limitations of measuring whole-plant traits non-destructively, we developed a low-cost and versatile approach that can non-invasively measure whole-plant physiology over time. As a proof of concept, we measured biomass accumulation and water use over time to directly quantify whole-plant water use efficiency. Plant water use efficiency (WUE), broadly defined as the ratio of biomass accumulation to water use via transpiration (Vadez et al., 2014), represents an important metric to optimize for improving plant resilience and agricultural sustainability. Various methods exist to quantify WUE, ranging from instantaneous measurements at the leaf-scale (Vadez et al., 2014), to long-term quantification at the canopy scale (Lambers and Oliveira, 2019). Because of the difficulty in measuring whole-plant WUE, it is common to extrapolate these data from leaf-level measurements or via proxies. For example, quantifying leaf gas exchange rates can provide a measure of both photosynthetic and transpiration rates. However, instantaneous leaf-level WUE data are often in disagreement with whole-plant measurements at longer timescales (Medrano et al., 2015). Leaf carbon isotope discrimination (CID) is often used to estimate whole-plant WUE as CID is strongly correlated with WUE in C_3_ species (Farquhar et al., 1982). However, the strength of this correlation is greatly reduced in C_4_ species, including important crops such as maize or sorghum (Henderson et al., 1998; Vadez et al., 2014). Moreover, quantifying CID is relatively slow and expensive (Vadez et al., 2014), thus necessitating simpler and more widely applicable methods.

In contrast to these methods, our approach, which uses hydroponic cultivation, allows for the direct and non-destructive measurement of whole-plant WUE through direct quantification of biomass accumulation and water use. We show that hydroponic cultivation results in WUE equal to that of soil-grown plants and then demonstrate the temporal and contextual versatility of our approach by measuring whole-plant WUE every two days on unstressed and osmotically-stressed seedlings. In doing so, we identified relationships between root zone acidification and aspects of photosynthetic efficiency on whole-plant growth and WUE over time. Our approach can be implemented using cheap, basic components and requires no specific technical expertise. It can be implemented for any non-aquatic, vascular plant species and can therefore contribute broadly to identifying and engineering superior crop germplasm.

## Results

We developed a simple, cheap, and modular system to grow and phenotype whole-plant traits non-destructively over time. A detailed, step-by-step protocol can be found online (protocols.io; dx.doi.org/10.17504/protocols.io.q26g7p6m1gwz/v1). Briefly, a hole is drilled into screw-on caps and a cut-off pipette tip of approximately equal diameter is inserted into the hole of the screw-on cap (Figure 1A). Soil is then inserted into the snugly-fit pipette tips and seeds are sown into the soil-filled tips. Plants are initially grown in “open” hydroponic conditions such that all samples are grown in a single reservoir whose water is exposed to the air (Figure 1B). Upon reaching a desired size, age, or developmental stage, tops are screwed onto tubes and the system becomes “closed” such that water loss from the tube is restricted to uptake by the plant (Figure 1C). By unscrewing the tops, the entire plant can be removed, weighed, and then screwed back onto the tube for continued growth (Figure 1C-D). When grown this way, whole-plant biomass accumulation and water use can be assayed repeatedly over time (Figure 2A-B). Moreover, the modularity of our approach allows for precise control of the root environment to investigate a variety of phenotypes, including the relationship of whole-plant biomass accumulation and water use in response to changing abiotic or biotic conditions.

**Figure 1:**
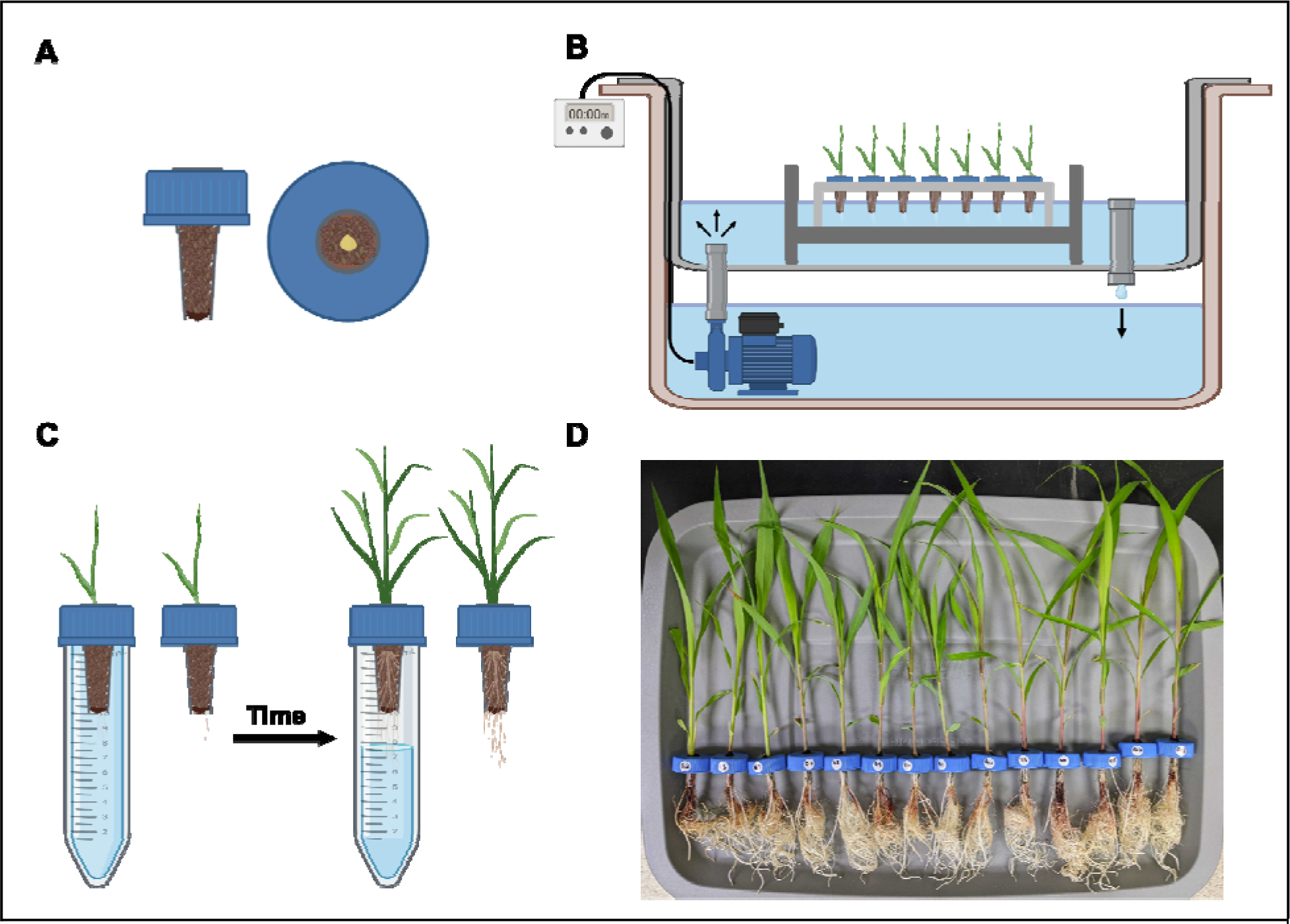
(A) Representation of a screw-on cap with a hole drilled into the top and a cut-off pipette tip inserted into the hole. Soil is inserted into the pipette tips and seeds are sown into the soil-filled tips. (B) Schematic of the recirculating ebb and flow hydroponic system used in this study. Water is pumped up from the bottom reservoir up to the top reservoir, where it rises up to the point of the drain pipe, and then falls back down into the bottom reservoir. Plants are placed in the top reservoir such that the water level rises up to the point of the soil-filled tips. The duration and time of day of irrigation events are controlled via a programmable timer connected to the pump. Samples are grown in these “open” hydroponic conditions until they reach a predetermined age or developmental stage. (C) Samples are then screwed onto tubes filled with nutrient solution, at which point the system is “closed”. When closed, water loss due to evaporation is prevented. After a designated amount of time, caps are unscrewed and the plants are lifted out of the tube. Tubes and caps with plants can be weighed to determine water use and biomass accumulation relative to previous timepoints, respectively. Caps with plants can then be screwed back onto tubes for continued growth in “closed” conditions. (D) Sorghum samples after 8 days of growth in tubes. Graphics for cartoon schematics in A-C are from BioRender.

**Figure 2:**
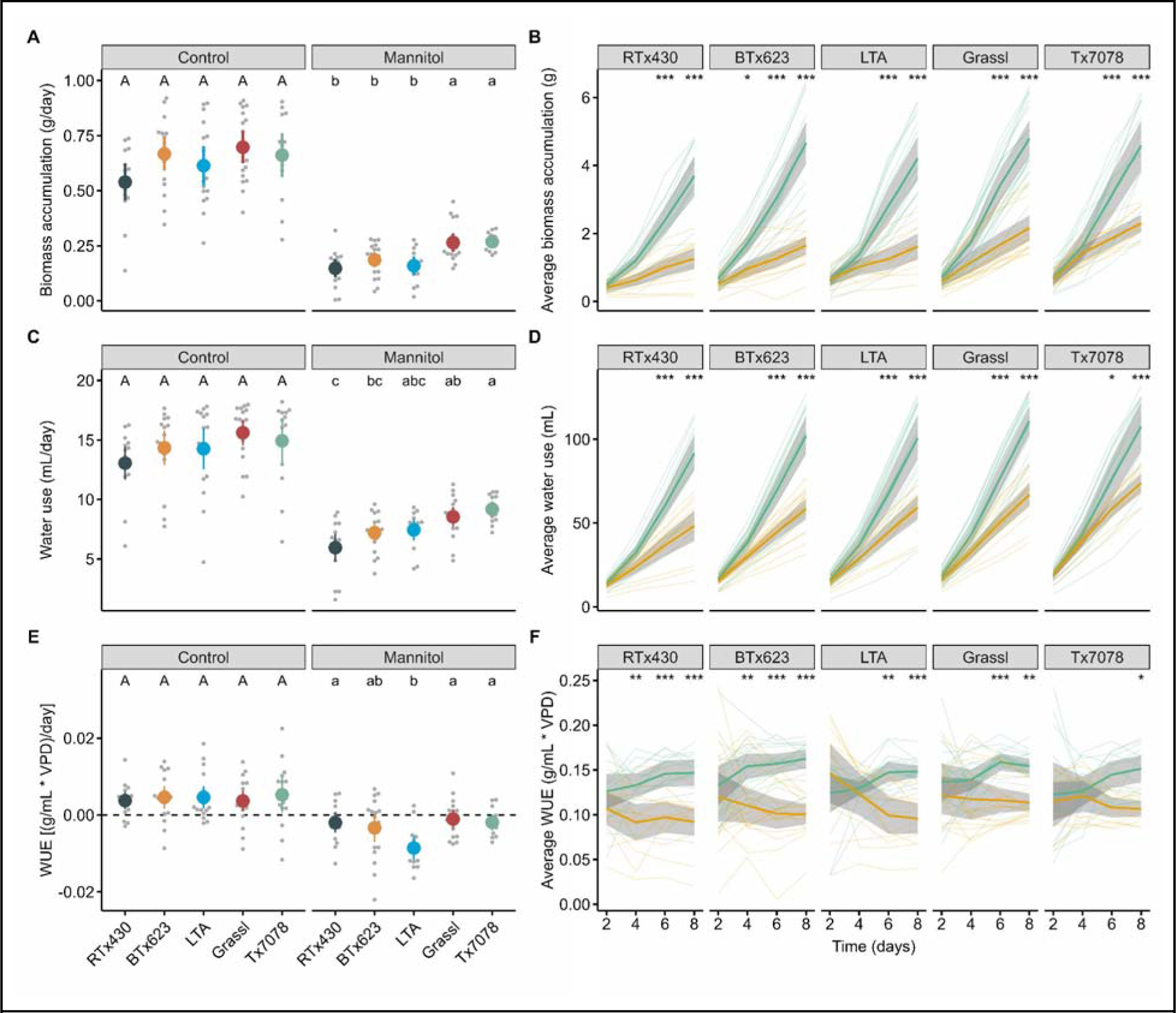
Average daily rate of biomass accumulation (**A**), water use (**C**), and WUE (**E**), as determined by linear regression (n = 12-18 per genotype, condition, and day; N = 4). Letters represent statistically different genotypes as determined by 1-way ANOVA (separately for control and mannitol treatments) followed by Tukey’s HSD test (alpha = 0.05). Cumulative biomass accumulation (**B**), water use (**D**), and WUE (**F**) of all genotypes over time in control (green) or mannitol-treated (orange) conditions. Thick solid lines in B, D, and F represent the average value per condition bounded by the 95% confidence interval. Thin lines in B, D, F represent individual samples. Asterisks indicate statistical significance at a given time point between mannitol and control samples as determined by 2-way ANOVA: * (p-value < 0.05), ** (p-value < 0.01), *** (p-value < 0.001). n, number of biological replicates per genotype and condition; N, number of independent experiments.

As a proof-of-concept of this approach, we sought to compare genotype-specific variation in biomass accumulation, water use, and whole-plant WUE of various sorghum accessions when grown either in soil or using our closed-system hydroponic approach. Sorghum (*Sorghum bicolor*) is a highly productive and relatively stress resistant C_4_ crop (Ananda et al., 2020; Khalifa and Eltahir, 2023). It is among the world’s most important cereal crops and is utilized for both food and bioenergy (Ananda et al., 2020; Khalifa and Eltahir, 2023). Genotype-specific variation exists in whole-plant WUE in soil-grown sorghum (Balota et al., 2008; Xin et al., 2008). We selected five sorghum genotypes (Table 1) to represent a diverse range of geographic origins, taxonomic races, variation in transpiration rates measured at the leaf-level (Balota et al., 2008), and variation in WUE measured at the whole-plant level (Balota et al., 2008, Xin et al., 2008).

**Table 1:**
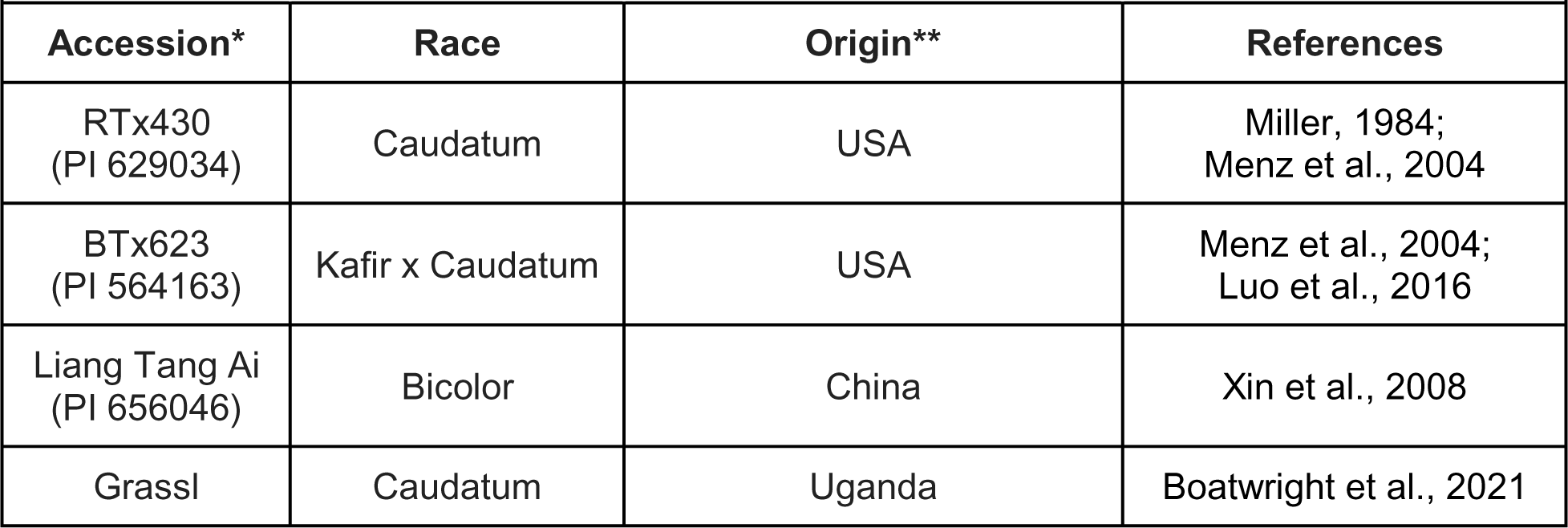
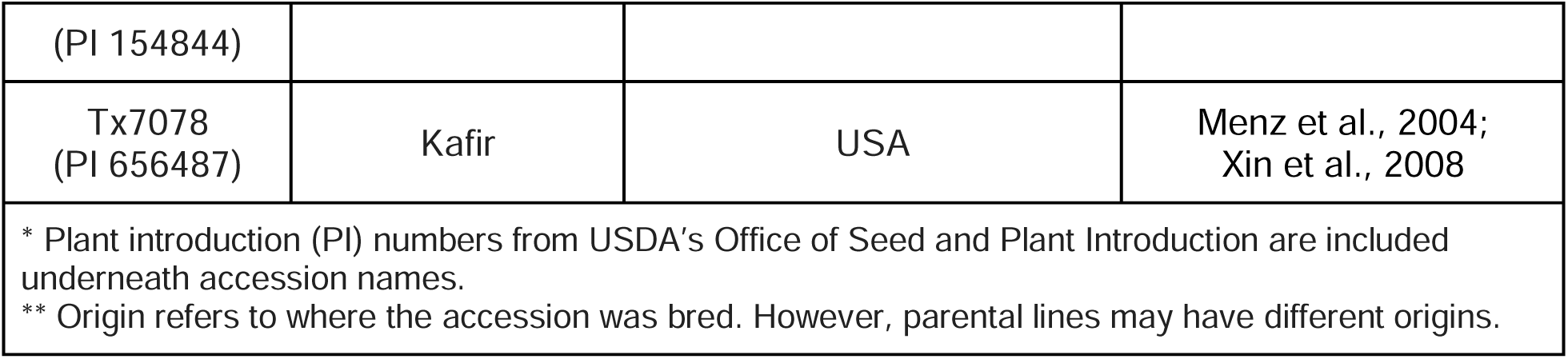
Description of sorghum accessions used in this study.

We first asked whether our method of quantifying WUE is consistent with established methods (Cernusak et al., 2006; Xin et al., 2009) by comparing the WUE of plants grown in our hydroponic system to plants grown in soil. Because most traditional soil-grown approaches to directly measure WUE are destructive, we compared end-point biomass accumulation and water use over the course of 8 days after reaching the third leaf stage in either hydroponic or soil-growth conditions. All genotypes accumulated more biomass and consumed more water when grown in soil compared to when grown hydroponically (Figure S2A-B). However, WUE after 8 days was indistinguishable between soil and hydroponic conditions for all genotypes (Figure S2C).

Next, we asked whether evaporative loss of water over time was negligible compared to plant water use, which is required to accurately infer plant water use from changes in tube weight over time. Water loss over time due to evaporation was measured in tubes from which whole plants were removed by cutting off roots and shoots to eliminate transpiration. Over the course of seven days, plant-less samples lost an average of 0.6 mL of water due to evaporation (Figure S3), equivalent to approximately 0.7% of average water use in plants grown in control conditions over the same period. Evaporative water loss in our hydroponic system was therefore determined to be negligible compared to transpirational water uptake.

Having validated our system, we next sought to compare the effect of osmotic stress on cumulative biomass accumulation, water use, and WUE across five sorghum accessions (Table 1). Specifically, we asked how these traits vary over time across each accession and whether genotype-specific differences in plant physiology manifest in unstressed or osmotically-stressed conditions. Plants were grown in tubes containing either nutrient solution alone or nutrient solution with 10mM mannitol to induce osmotic stress. Osmotic stress resulted in decreased biomass accumulation, water use, and WUE in all genotypes, compared to control conditions (Figure 2). There were no genotype-specific differences in the rates of biomass accumulation, water use, or WUE under control conditions (Figure 2A,C,E). However, genotype-specific differences were observed in both the time and extent to which mannitol stress impacted growth and WUE. For example, Grassl and Tx7078 accumulated biomass at a greater rate than the other genotypes when grown in mannitol (Figure 2A). Additionally, biomass accumulation in osmotically stressed BTx623 was reduced compared to control conditions after 4 days, which was earlier than all other genotypes (Figure 2B). Despite this temporal difference, the overall rate of biomass accumulation in BTx623 under osmotic stress was similar to that of osmotically-stressed RTx430 and LTA (Figure 2A). Mannitol stress resulted in the greatest reduction in water use rates in RTx430 and the smallest decrease in Tx7078 (Figure 2C). However, there were no genotype-specific temporal differences in average water use rates under osmotic stress compared to controls (Figure 2D). With regards to WUE rates, stress-induced reductions were most pronounced in LTA (Figure 2E). This greater reduction in WUE is in contrast to the time at which WUE levels of osmotically-stressed plants dropped below control levels. Specifically, WUE of RTx430 and BTx623 plants grown in mannitol was decreased compared to control by Day 4, while osmotic stress did not reduceTx7078 WUE until Day 8, later than all other genotypes (Figure 2F). Collectively, these results suggest that the onset of osmotic stress does not always mirror its magnitude on plant physiology over time. Additionally, these results indicate that Grassl and Tx7078 are more tolerant to osmotic stress than RTx430 and that LTA is less efficient in using water compared to other accessions when osmotically stressed at the third-leaf stage.

To better understand how osmotic stress affects the relationship between biomass accumulation and water use, and to identify which underlying component is a stronger predictor of changes in WUE, we regressed biomass accumulation by water use (Figure S4A), WUE by biomass accumulation (Figure S4B) and WUE by water use (Figure S4C). There was a strong positive correlation between biomass accumulation and water use for all genotypes in both control and mannitol-treated samples, though this relationship was stronger in control conditions (Figure S4A). A positive correlation was also found between WUE and biomass accumulation for all genotypes under both control and osmotically-stressed conditions. However, the correlation was stronger for all genotypes when exposed to osmotic stress (Figure S4B). Conversely, WUE could only be explained in part by water use in certain genotypes and under certain conditions (Figure S4C). In LTA, for example, WUE increased linearly with water use only under osmotic stress, whereas there was no relationship between WUE and water use in either condition for Tx7078 (Figure S4C). For all genotypes in which a positive correlation was found between both biomass accumulation and water use with WUE, biomass accumulation was the stronger predictor for increases in WUE (Figure S4B-C). These results suggest that mannitol-induced osmotic stress reduces water use efficiency and that in both stressed and unstressed conditions, biomass accumulation is a stronger determinant of WUE than water use.

Having observed genotype-specific differences in biomass accumulation, water use, and WUE rates specifically in osmotically-stressed conditions, we next sought to identify possible mechanisms for this context-specific difference in plant growth and physiology. Osmotic stress can impair photosynthetic performance (Grzesiak et al., 2006; Chen et al., 2011). We therefore asked whether genotype-specific differences in biomass accumulation, water use, and WUE under mannitol stress could be explained by differences in photosynthetic efficiency.

Photosystem II (PSII) quantum efficiency (F_V_/F_M_) of the third true leaf decreased to the greatest extent in Tx7078 and RTx430 plants in response to osmotic stress (Figure 3A). Similar to the effects of osmotic stress on biomass accumulation and water use, the time at which F_V_/F_M_ levels dropped compared to control varied by genotype (Figure 3B). Specifically, F_V_/F_M_in mannitol-grown LTA plants was reduced relative to control levels by Day 4, which was earlier than all other genotypes (Figure 3B) and further demonstrates that the onset of stress does not always correlate with its magnitude. These results suggest that photosynthetic capacity at the third-leaf stage was not strongly associated with plant growth or WUE under osmotic stress, as Grassl and Tx7078 plants had equal rates of biomass accumulation and WUE under osmotic stress despite differences in the effect of osmotic stress on their photosynthetic efficiency.

**Figure 3:**
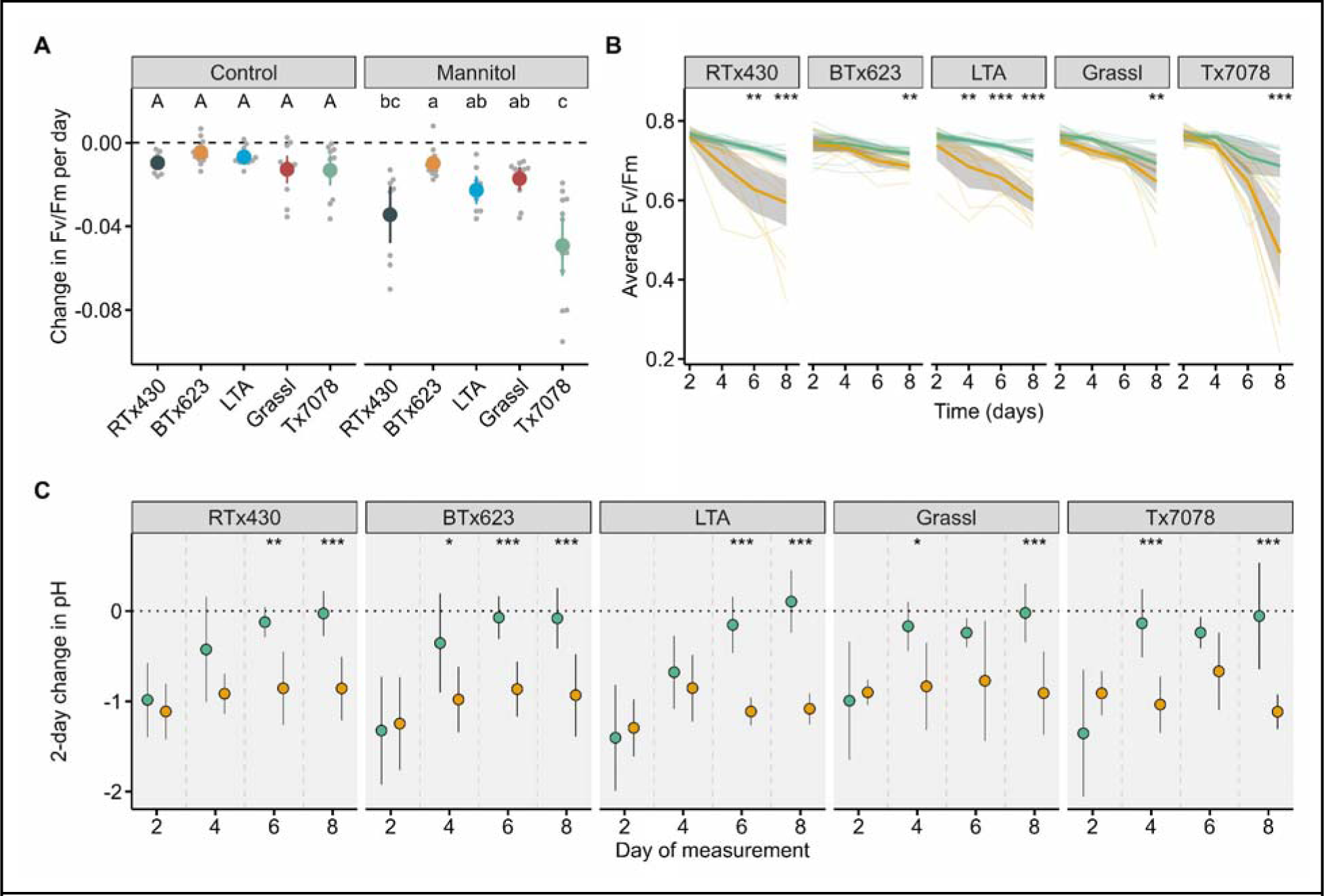
(**A**) Average rate of daily change in F_V_/F_M_ as determined by linear regression (n = 9-18 per genotype, condition, and day; N = 3-4). Letters represent statistically different genotypes as determined by 1-way ANOVA (separately for control and mannitol treatments) followed by Tukey’s HSD test (alpha = 0.05). (B) Average F_V_/F_M_ over time of the third true leaf of samples grown in control (green) or mannitol (orange) conditions. Thick solid lines represent the average value per condition bounded by the 95% confidence interval. Thin lines represent individual samples. (C) Average 2-day change in pH in the growth media in control (green) or mannitol (orange) conditions (n = 7-18 per genotype, condition, and day; N = 3-4). Note that any solution remaining in the tubes after every 2-day period was discarded and replaced by fresh nutrient solution to ensure sufficient water for plant growth for the subsequent two days. Error bars represent the standard deviation. Asterisks in B and C indicate statistical significance at a given time point between mannitol and control samples as determined by 2-way ANOVA: * (p-value < 0.05), ** (p-value < 0.01), *** (p-value < 0.001). n, number of biological replicates per genotype and condition; N, number of independent experiments.

We next asked if there were any differences between the accessions in their root physiology in response to osmotic stress. By exuding a diverse set of compounds from their roots, plants actively modify their root zone to facilitate nutrient uptake and in response to abiotic stress, including mannitol-induced osmotic stress (Marschner et al., 1986; Falhof et al., 2016; Darko et al., 2019). One of the ways in which plants optimize their rhizosphere properties is via soil acidification (Marschner et al., 1986; Ehrenfeld et al., 2005). Growing plants in closed tubes makes it possible to directly and easily measure rhizosphere acidification. We therefore asked whether and when there were any genotype-specific differences in pH change of the nutrient solution after every two days (delta-pH). The pH of nutrient solution with and without mannitol-supplementation was measured before being added to tubes, and the remaining solution in tubes after two days of plant growth was then measured again (Figure 3C, Supplemental File 1). Any solution remaining in the tubes after every two days was then discarded and fresh nutrient solution was added to ensure sufficient water for plant growth for the subsequent two days. After two days of growth in closed tubes, there was no effect of osmotic stress on delta-pH in any genotype (Figure 3C). However, the root zone was acidified more in response to osmotic stress by Day 4 in BTx623, Grassl, and Tx7078 plants and by Day 6 in RTx430 and LTA plants (Figure 3C). While control plants of all genotypes generally ceased acidifying their nutrient solution by Day 8 at the latest, osmotic stress resulted in continuous root zone acidification throughout the experiment for all genotypes (Figure 3C, Supplemental File 1). These results suggest that prolonged root zone acidification is an adaptive response conserved among sorghum accessions in response to osmotic stress at the seedling stage.

We have demonstrated how non-destructive, whole-plant phenotyping can be employed using simple and readily available components. We showcase the versatility of our approach by directly measuring multiple traits across a diverse set of sorghum accessions in unstressed and osmotically-stressed conditions. Collectively, we illustrate how this method can identify genotype-specific temporal dynamics in plant growth across multiple organs and physiological processes.

## Discussion

Plant phenotypes are dynamic manifestations of genetics, environmental conditions, and developmental stage. Phenotypic plasticity is evidenced by changes in growth of both above- and below-ground tissues. Non-destructive, whole-plant phenotyping facilitates quantification of dynamic growth processes which are more challenging to identify via destructive approaches and is therefore crucial for improving our understanding of plant growth and for engineering more resilient and resource-efficient crops. However, noninvasive, whole-plant phenotyping approaches are often expensive and technically complex. To address these limitations, we developed a cost-effective, simple, and modular approach that can measure whole-plant physiology over time. Unlike soil-grown conditions, hydroponic cultivation allows us to physically isolate roots and return them to their growth environment non-destructively. In doing so, entire plant biomass can be measured directly and repeatedly. Additionally, if each plant is grown physically separated from other plants and evaporation is prevented, plant water use can also be measured directly. These capabilities alone are of practical value for the study of plant growth and development given that these traits are not readily quantifiable in a direct, non-destructive manner for soil-grown plants. Moreover, quantification of these traits over time allows for direct measurements of whole-plant WUE over time. Previous studies have demonstrated creative approaches to quantify WUE non-destructively (Fahlgren et al., 2015; Fletcher et al., 2018). However, these methods either ignore root biomass contributions (Fahlgren et al., 2015), or do not correlate well with WUE values from harvested soil-grown plants (Fletcher et al., 2018).

Comparing our hydroponic system to potted soil, we observed that plant growth and resource usage were strongly influenced by abiotic conditions. This was made clear by the greater biomass accumulation and water use of soil-grown plants compared to those grown hydroponically. Interestingly, WUE was uniform across growth conditions. This suggests that biomass accumulation at this stage of development necessitates a proportionally equal amount of water use, regardless of absolute growth rates. What might explain slower growth rates in the hydroponic system compared to potted soil? The hydroponic growth conditions used in this study were chosen to be suitable for a diverse range of genotypes at a specific developmental stage, rather than being designed to optimize growth. However, irrigation scheduling, nutrient composition, and abiotic conditions for hydroponic growth could all be refined to optimize a given metric, as desired.

We chose to examine the five varieties of sorghum (Table 1) because, previously, genotype-specific differences in whole-plant WUE have been observed in soil-grown sorghum (Mortlock and Hammer, 1999; Xin et al., 2008; Xin et al., 2009). We specifically focused on identifying genotype-specific variation at the seedling stage as sorghum biomass accumulation at the time of harvest is most affected by drought-induced osmotic stress when stress occurs at the seedling stage (Mastrorilli et al., 1999), similar to the heightened sensitivity of many crop species to stresses at early stages in their development, such as maize (Kang et al., 2000; Ge et al., 2012), rice (Zhao et al., 2014), and tomato (Sivakumar et al., 2020). Under non-stressed conditions, LTA previously exhibited greater whole-plant WUE than Tx7078 when grown in soil, but equal WUE to BTx623 and RTx430 (Xin et al., 2008; Xin et al., 2009). That we did not observe genotype-specific differences in WUE in either soil- or hydroponically-grown sorghum in control conditions could reflect the different ages and developmental stages of the plants being assayed. We measured plant growth starting at the third leaf stage, corresponding to plants which were approximately 2-weeks old, whereas plants were approximately 4-6 weeks old (corresponding to the 8th-leaf stage) in Xin et al., 2008 and Xin et al., 2009. The differences in WUE we observed compared to previous reports further support the idea that WUE varies by developmental stage (Fahlgren et al., 2015). We did, however, observe genotype-specific differences in WUE when plants were grown under osmotic stress. In light of the previously-identified superiority of LTA in terms of its WUE when grown in soil under non-stressed conditions, it was somewhat surprising that LTA was most sensitive to osmotic stress in terms of its WUE. While WUE of many species, including sorghum (Xie and Su, 2012), increases in response to drought stress in soil-grown conditions (Peters et al., 2018), mannitol-induced osmotic stress decreases WUE in multiple species (Cha-Um and Kirdmanee, 2008; Darko et al., 2019), similar to what we observed with sorghum in this study. Collectively, these findings highlight the importance of considering multiple developmental stages and abiotic conditions when studying plant resilience.

Because WUE is underpinned by both biomass accumulation and water use, much work has been done to identify which of these components should be optimized to improve crop WUE. Whether WUE is even a desirable agronomic trait is actually a contentious matter (Condon et al., 2004; Blum, 2009). This is because, in many crop species, increased WUE is usually achieved through reduced stomatal conductance, which in turn results in decreased carbon fixation and lower yields (Blum, 2009; Vadez et al., 2014). Other work, however, has demonstrated that higher WUE via reduced stomatal conductance does not necessitate a yield penalty if there is a concomitant increase in photosynthetic rates (Vadez et al., 2014). These contradictory findings suggest that the contributions of growth and water use to WUE are both complex and are underpinned by both genotypic and environmental factors. In all such cases, however, it is generally agreed that increasing, rather than decreasing soil water use, is a key breeding strategy for improving both crop yield and potentially WUE (Blum, 2009; Condon, 2020). This is because increased plant water use would support more carbon capture and reduce the fraction of soil water lost to evaporation. Moreover, increased root growth could facilitate water capture from deeper in the soil profile where water is less prone to evaporative losses (Condon, 2020). These findings suggest that breeders should aim to identify genotypes which maximize water use and biomass accumulation, as these traits should support both high yields and plant resilience during water stress (Blum, 2009; Condon, 2020). This idea is supported by the findings in this study, given that WUE positively correlated with both increased biomass accumulation and water use and that these relationships were stronger under osmotic stress. Moreover, biomass accumulation, rather than water use, was a stronger predictor of WUE in all genotypes, which has previously been observed in soil-grown sorghum (Xin et al., 2009). These results further support the idea that breeders should aim to identify the fastest growing lines to both optimize yields and WUE while bearing in mind the complex interactions of WUE with plant genetics, developmental stage, and abiotic conditions.

The ability to track plant growth over time non-destructively can reveal biological insights that could be missed through endpoint analysis alone (Fahlgren et al., 2015). For example, average WUE of mannitol-treated RTx430 and BTx623 plants was already lower than control plants by Day 4, yet F_V_/F_M_ in those same plants was not yet distinguishable compared to control, suggesting that WUE in sorghum is negatively affected by osmotic stress before photosystem II quantum efficiency. Also, because biomass accumulation was already reduced by Day 4 in mannitol-treated BTx623 plants compared to control, something other than a reduction in photosynthetic efficiency was impairing growth rates in that accession. End-point analysis would not have revealed how osmotic stress impacted growth and photosynthetic efficiency differently and at different times among the genotypes. Non-destructive phenotyping also brought to light that stress onset does not necessarily correlate with stress magnitude, as evidenced by the late, but severe drop in Fv/Fm in Tx7078 plants compared to the other genotypes. Previous work utilizing non-destructive phenotyping techniques has also identified a disconnect between stress onset and magnitude in Arabidopsis (Awlia et la., 2016). Collectively, these observations further support the value of non-destructive approaches to capturing the dynamic nature of plant phenotypes across time and environments (Granier and Vile, 2014).

Plants actively modify the soil properties of their growing environment to optimize growth by improving nutrient uptake and mobilization and to regulate soil microbial dynamics (Marschner et al., 1986; Shen et al., 2013; Falhof et al., 2016). Depending on the biotic and abiotic conditions in the soil, roots can exude sugars, amino acids, enzymes, organic acids, and phytosiderophores to optimize conditions for symbiotic rhizospheric microorganisms (Ryan et al., 2009) or for direct nutrient uptake (Marschner et al., 1986; Shen et al., 2013). The non-destructive nature of our closed-system hydroponic approach allowed us to more readily identify a connection between root-zone acidification and biomass accumulation than had we grown plants in soil for destructive end-point analyses. For example, after two days of growth in closed tubes, both control and mannitol-treated plants of all genotypes acidified their root zones equally. At this time point, there was also no difference in total biomass accumulation between control and mannitol-treated samples in any genotype. While root zone acidification gradually decreased over time in control samples of all genotypes, it remained fairly constant for mannitol-treated plants. That mannitol-treated plants continued to acidify their root zone throughout the experiment suggests a greater and prolonged investment of root exudates to actively modify their root environment. This could be because mannitol-stress can impair uptake and translocation of phosphorus (Resnik, 1970) and potassium (Slama et al., 2007). Interestingly, the average pH after every two days of growth in mannitol was 5.5-5.8 (Supplemental File 1), which is a range that favors plant uptake of all essential nutrients (Reed, 1996; Pennisi and Thomas, 2009). Thus, root zone acidification in response to mannitol stress could be an adaptive response to improve mineral nutrient uptake and mobility.

Various aspects of the dynamic effects of root-zone conditions on plant growth remain poorly understood (Kardol et al., 2013). Additionally, many plant responses to environmental conditions are age-dependent (Bond, 2000; Panter and Jones, 2002). Thus, while hydroponic cultivation is an established approach in plant biology research (Asao, 2012), the isolation and modularity of the root zone environment afforded by our approach open the door to additional avenues of experimentation for directly studying dynamic, and age-dependent processes. For example, researchers could finely control the characteristics of the solution in each tube to investigate temporal effects of mineral nutrition or biotic stress on whole-plant development and physiology and to analyze how these characteristics differentially affect shoot vs. root growth.

Our approach also makes it easier to investigate how changes in root architecture and morphology over time influence plant growth. While currently our approach does not distinguish shoot from root biomass, it could easily be paired with imaging-based phenotyping technologies such as PhenoBox (Czedik-Eysenberg et al., 2018) or PlantCV (Gehan et al, 2017) to quantify both components separately. Future studies which incorporate such imaging technologies could directly quantify the relationship between root zone acidification and root growth characteristics. Some plant species require more oxygen in their root zone than others (Narsai et al., 2011; Asao, 2012). Supplementing hydroponic nutrient solution with hydrogen peroxide can increase the availability of dissolved oxygen in the growing media and can increase nutrient uptake and chlorophyll content in various species (Butcher et al., 2017; Liu et al, 2022). While suitable concentrations of hydrogen peroxide would need to be established beforehand to avoid phytotoxicity, its use in the nutrient solution would not require any physical modifications to the closed-hydroponic design. Additionally, tubes could be opened more frequently than every two days, to further increase root zone oxygenation. While 50mL tubes were sufficiently large for sorghum plants at the 3rd leaf stage, larger tubes could be used for either older or larger species, as required.

Efforts to identify and engineer more resource-efficient and resilient crops will benefit from phenotyping approaches which can noninvasively quantify whole-plant growth and physiology over time. Our approach allows for inherently greater throughput than traditional approaches and can, in its entirety, be implemented using cheap and readily available components. Its versatility for studying various aspects of plant growth and physiology make it a valuable method for plant scientists as a whole in studying plant growth and for identifying superior and more resilient crops.

## Materials and Methods

A detailed, step-by-step protocol can be found at protocols.io; dx.doi.org/10.17504/protocols.io.q26g7p6m1gwz/v1.

An R notebook containing scripts for visualizing and analyzing data from these experiments can be found at https://rpubs.com/danny_ginzburg/1143647.

### Closed-system hydroponic growth apparatus construction and seeding

Screw-on tops of 50mL Falcon tubes were used to both physically support growing plants and to secure individual tubes to prevent evaporative water loss. To physically support growing plants, holes were drilled into the center of 50mL Falcon tube caps using a 21/64 inch drill bit. 1250 µL pipette tips were cut ∼3 cm from the tip (narrower) end and inserted into the pre-drilled Falcon tube caps such that the tips fit snugly into the holes in the caps and rested almost flush with the top of the cap. Rapid-Rooter starter plugs (General Hydroponics, Santa Rosa, CA, USA) were cut into rectangular slices of ∼0.5cm length and ∼0.9cm width and were inserted into the top of each tip and then gently pushed to the bottom. To prevent unnatural water retention and pressure buildup in the filled tips, pinholes were created in the tips ∼25% from the bottom using a needle. Caps with tips were then filled with fresh Pro Mix Bio-Fungicide potting soil (Premier Tech Horticulture, Quakertown, PA, USA), placed in tube racks, and left to soak overnight in tap water which reached the bottom of the tops but did not submerse the caps. The next day, individual sorghum seeds were planted into soil-filled tips. While still in racks, all samples were then placed into an in-house built ebb & flow (flood & drain) hydroponic system.

Briefly, an ebb & flow system utilizes a submersible pump to recirculate nutrient-rich water between two or more reservoirs. Plants are placed within one of the reservoirs and grown hydroponically. The ebb & flow system used in this study consisted of two, 18 gallon Rubbermaid Roughneck plastic totes (reservoirs) (Rubbermaid, Atlanta, GA, USA), one stacked on top of the other (Figure S1A). Two, ¾ inch diameter holes were drilled in opposite corners of the top reservoir. A ¾ inch threaded and barbed bulkhead fitting (Figure S1B) (Botanicare, Vancouver, WA, USA) was inserted from above into one of the drilled holes and fastened from below with a rubber washer. Water in an ebb and flow system needs to reach, but not submerge, the seedlings in the upper reservoir. Thus, to increase the irrigation height the water reaches in the upper reservoir before draining back down, a threaded height extender (Figure S1B) was then screwed into this first hole from above. The second hole which was drilled into the upper reservoir was then fitted with a threaded, plastic debris screen (Figure S1B) to allow water to gently pump up into the top reservoir. In the bottom reservoir sat an 8W, 800 liter/hour submersible pump (VIVOSUN, Ontario, CA, USA) connected via PVC plastic tubing to the barbed end of the bulkhead fitting (Figure S1A). When turned on, water is pumped up to the top reservoir from below and rises up to the level of extender before draining back down to the bottom reservoir. In order for water to constantly recirculate during irrigation, sufficient water must be added to the ebb & flow system to allow for filling the top reservoir up to the draining height and to ensure enough water is in the bottom reservoir to be pumped back up. To control the duration and number of irrigation events each day, the pump was connected to a digital programmable timer (BN-LINK, Santa Fe Springs, CA, USA).

### Pre-phenotyping hydroponic growth conditions

The drainage height of the ebb & flow system was optimized such that, during irrigation, the bottom ∼2-3cm of the cut pipette tips would be submerged in water (Figure 1B). Before seedling emergence, samples were irrigated thrice daily with tap water for 5 minutes every 8 hours using a programmable timer as described above. Upon root emergence (root tips growing below the bottom of the pipette tip) of 50% of the samples, seedlings were irrigated every hour with a single, 15-minute irrigation event using a programmable timer. At this point, Peters Professional General Purpose 20-20-20 soluble fertilizer (ICL Specialty Fertilizers, Tel Aviv, Israel) was directly added to the lower water reservoir using a 50mL plastic scoop up to an electrical conductivity (EC) of 1200 mS/cm (∼1.25g fertilizer/liter water) while the water was being circulated. EC was measured with a Hanna Edge conductivity meter (Hanna Instruments, Rhode Island, USA). Upon root emergence of all samples, irrigation duration was increased to 45-minute events every hour followed by 15 minutes of no irrigation to increase root zone oxygenation. The reservoir was emptied and refilled with fresh nutrient solution every 3-4 days. All experiments were conducted in a greenhouse with supplemental lighting from high pressure sodium lamps from 7am-7pm daily.

### Transitioning samples from open-to closed-hydroponic conditions

Once 50% of samples reached the 3rd true-leaf stage, a bulk nutrient solution was prepared such that 40mL of solution could be allocated to each sample in 50mL Falcon tubes. Nutrient solution in 50mL tubes consisted of 0.4g/L Hoagland’s No. 2 basal salt mixture (Sigma Aldrich Chemical Co. St. Louis, MO, USA) supplemented with ∼0.5g/L Professional General Purpose 20-20-20 fertilizer up to an EC of 1200 mS/cm. This bulk nutrient solution was then split into two, and mannitol was added into one of the solutions up to a concentration of 10mM. pH of both control and mannitol-supplemented solutions was then recorded. 50 mL tubes were then filled with 40mL of either control or mannitol-supplemented nutrient solution. Tubes were wrapped in aluminum foil to prevent algal growth. Tubes with solution, but no caps, were then weighed. Caps containing seedlings were then brought into the lab and were firmly screwed onto individual tubes. Tubes with seedlings screwed on were then weighed to determine initial seedling weight. If the roots of any sample did not reach the nutrient solution when fully closed into the cap, additional solution was added and the tube (with and without cap) was reweighed. Samples were evenly spaced into 50mL tube racks then returned to the greenhouse.

### Quantifying biomass accumulation, water use, and change in media pH

Every two days, fresh nutrient solution was prepared and pH was measured as described above for both control and 10mM-treated samples. Seedlings and caps were unscrewed from their tubes and placed back into the ebb & flow system (operating with constant irrigation) to keep the roots hydrated while tubes were weighed. 50mL tube caps (without holes) were screwed onto each tube to prevent evaporation. Tubes were brought back into the lab and were individually weighed (without caps). pH of the solution in each tube was also recorded at this time. Tubes were emptied of their solution and refilled with new control or 10mM mannitol-supplemented solution up to 40mL as described above. Tubes without caps were then weighed again and caps were screwed back on to prevent evaporation. Seedlings were brought into the lab, individually weighed, and then screwed back onto their corresponding tubes before being returned to the greenhouse.

### Chlorophyll fluorescence quantification

In the greenhouse, seedlings were gently laid on their side and dark-adaptation clips were placed onto the middle of the 3rd true leaf of each sample for 30 minutes. After dark-adaptation, F_V_/F_M_ of the 3rd true leaf was recorded using a chlorophyll fluorometer (OS30p+, Opti-Sciences, Inc. Hudson, New Hampshire). Specifically, minimum fluorescence (F_0_) was measured after application of a weak modulated pulse of 0.1 μmol m^−2^ s^−1^ PPFD to the sampled region. A 1-second saturating pulse of 6,000 μmol m^−2^ s^−1^ PPFD was subsequently applied to the same region to determine maximum fluorescence (F_M_). F_V_/F_M_ was calculated as (F_M_−F_0_)/F_M_. F_V_/F_M_ measurements were always made between 2-3pm, as chlorophyll fluorescence parameters are under circadian regulation (Dodd et al., 2014). Samples were then placed upright back into tube racks.

### Determination of vapor pressure deficit

In addition to genetic and developmental factors, plant transpiration rates are driven by vapor pressure deficit (VPD) (Mortlock and Hammer, 1999; Vadez et al., 2014; Grossiord et al., 2020), which is the difference between ambient vapor pressure and vapor pressure in saturated air (Grossiord et al., 2020). To facilitate comparisons of WUE across time and experiments, we therefore normalized WUE at each timepoint by the average cumulative daytime VPD (Sinclair, 2012; Vadez et al., 2014). Temperature and humidity were measured continuously with a Govee H5075 thermometer-hygrometer (Govee Moments Trading Limited, Hong Kong) and were averaged over 15-minute intervals. VPD was derived from temperature and RH values using the following equations from Murray (1967):

1. Saturation vapor pressure (SVP) in kPA = 0.61078 * e^(T/(T^ ^+^ ^237.3)^ ^x^ ^17.2694)^ where T is temperature in degrees Celsius
2. Actual vapor pressure (AVP) = (SVP*RH)/100 where RH is percent relative humidity
3. VPD = SVP - AVP

Daytime VPD was calculated as the average VPD during the hours when supplemental lighting was provided in the greenhouse, namely 7am-7pm.

### Determination of evaporative loss in closed-hydroponic system

Plants were grown hydroponically in soil-filled pipette tips as described above. At the 3rd leaf stage, roots and shoots were cut from the bottom of the tips and the base of the cap, respectively to prevent plant water uptake. An equal number of tubes as caps were filled with 1200 EC nutrient solution and weighed, as described above. Caps from which roots and shoots were cut off were then screwed onto nutrient-filled tubes. Samples were placed in the same greenhouse as described above. Over the course of seven days, caps were removed from tubes and tubes were weighed to determine average daily and cumulative water loss due to evaporation. Caps were then screwed back on after weighing. This experiment was repeated three times. Average 7-day water use on control plants was derived from the linear regression model of water use as a function of time. Cumulative water loss of plant-less samples was then divided by the computed 7-day plant water use to represent the percentage of plant water use.

### Determination of water use efficiency from soil-grown plants

Seeds were planted in pots filled with an equal amount, by weight, of Pro Mix Bio-Fungicide potting soil. Soil-grown plants were grown in the same greenhouse as hydroponically-grown plants, as described above. When 50% of the samples reached the 3rd true leaf stage, at least three samples per genotype were harvested to determine average total biomass per genotype (initial biomass). Samples harvested at the 3rd true leaf stage were visually of similar size to those which were not yet harvested. For the remaining samples, pots were watered up to the point of soil saturation and were then weighed. The soil surface and drainage holes at the bottom of each pot were then covered with aluminum foil to prevent evaporative water loss. After 8 days of growth, the aluminum foil was removed and pots with plants were weighed. Plants were then harvested to determine final, whole-plant biomass. Eight-day biomass accumulation was calculated as (final biomass - initial biomass). Water use was calculated as (initial pot weight - average initial biomass) - (final pot weight - final whole-plant biomass). Soil-grown WUE was normalized by average cumulative daytime VPD, as described above.

### Statistical Analyses

Linear regressions and their statistical comparisons were calculated using the lm() and lstrends() functions, respectively, from the ‘lsmeans’ package (Lenth, 2016) in R version 4.2.2. Differences in control vs mannitol averages at each time point were calculated by two-way ANOVA followed by Tukey’s HSD test (p-value < 0.05) using the aov() function from the Base R ‘stats’ package (R Core Team, 2013). Slopes and r-squared values from linear regressions were calculated using the stat_regline_equation() function from the ‘ggpubr’ package (Kassambara, 2023). Comparisons of biomass accumulation, water use, and WUE between soil- and hydroponically-grown samples were calculated by two-way ANOVA followed by Tukey’s HSD test (p-value < 0.05) using the lsmeans() function.

---

**Supplemental File 1:** Initial pH, final pH, and delta-pH for all genotypes, conditions, and days of measurement.

## Acknowledgements

We thank I. Villa for help with drilling holes in both the ebb and flow system and all the screw-on caps, G. Materassi-Shultz for plant growth facility support, S. Mangat for help collecting data, members of the Rhee lab for helpful discussions, and the USDA National Plant Germplasm System for providing sorghum seeds. This work was done on the ancestral land of the Muwekma Ohlone Tribe, which was and continues to be of great importance to the Ohlone people.

## Author contributions

D.N.G. conceived of and designed the closed-hydroponic apparatus. D.N.G. and S.Y. R. conceived of the experimental design. D.N.G. and J.A.C. performed the growth experiments. J.A.C. and S.Y.R. provided intellectual contribution and input into manuscript organization. D.N.G. wrote the manuscript and S.Y.R. edited the manuscript. S.Y.R. advised D.N.G. and J.A.C.

## Data availability

All study data are included in the main text and supporting information.

---

**Figure S1:**
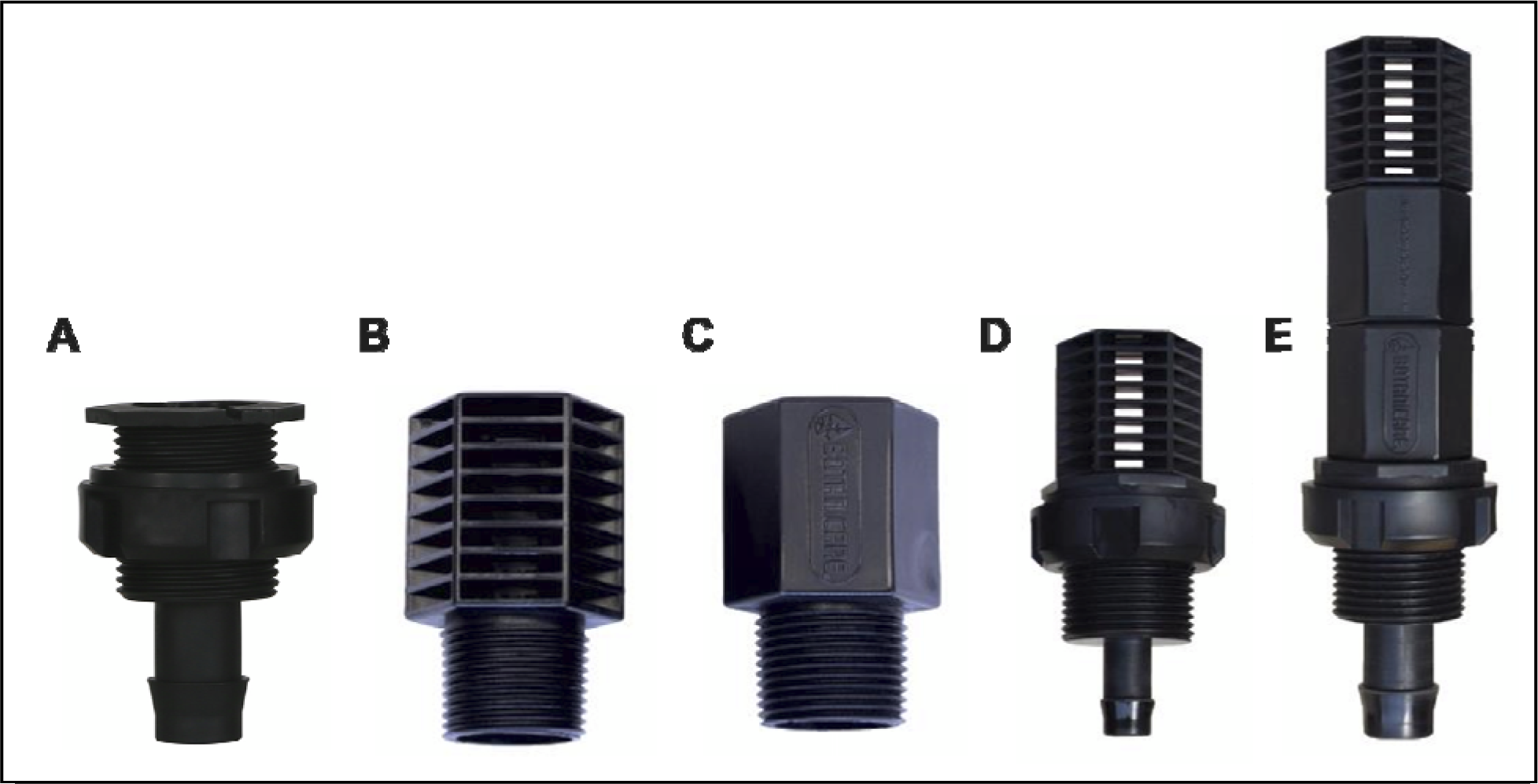
Images of Botanicare fittings used in the ebb and flow system. (A) threaded bulkhead fitting which is inserted into both holes that are drilled into the upper reservoir; (B) threaded plastic debris screen; (C) threaded height extender; (D) debris screen screwed onto bulkhead fitting for water inflow into the upper reservoir; (E) debris screen screwed onto two height extenders for water flowing out of the upper reservoir and back down into the lower reservoir. Images of Botanicare fittings are from https://www.hawthornegc.com.

**Figure S2:**
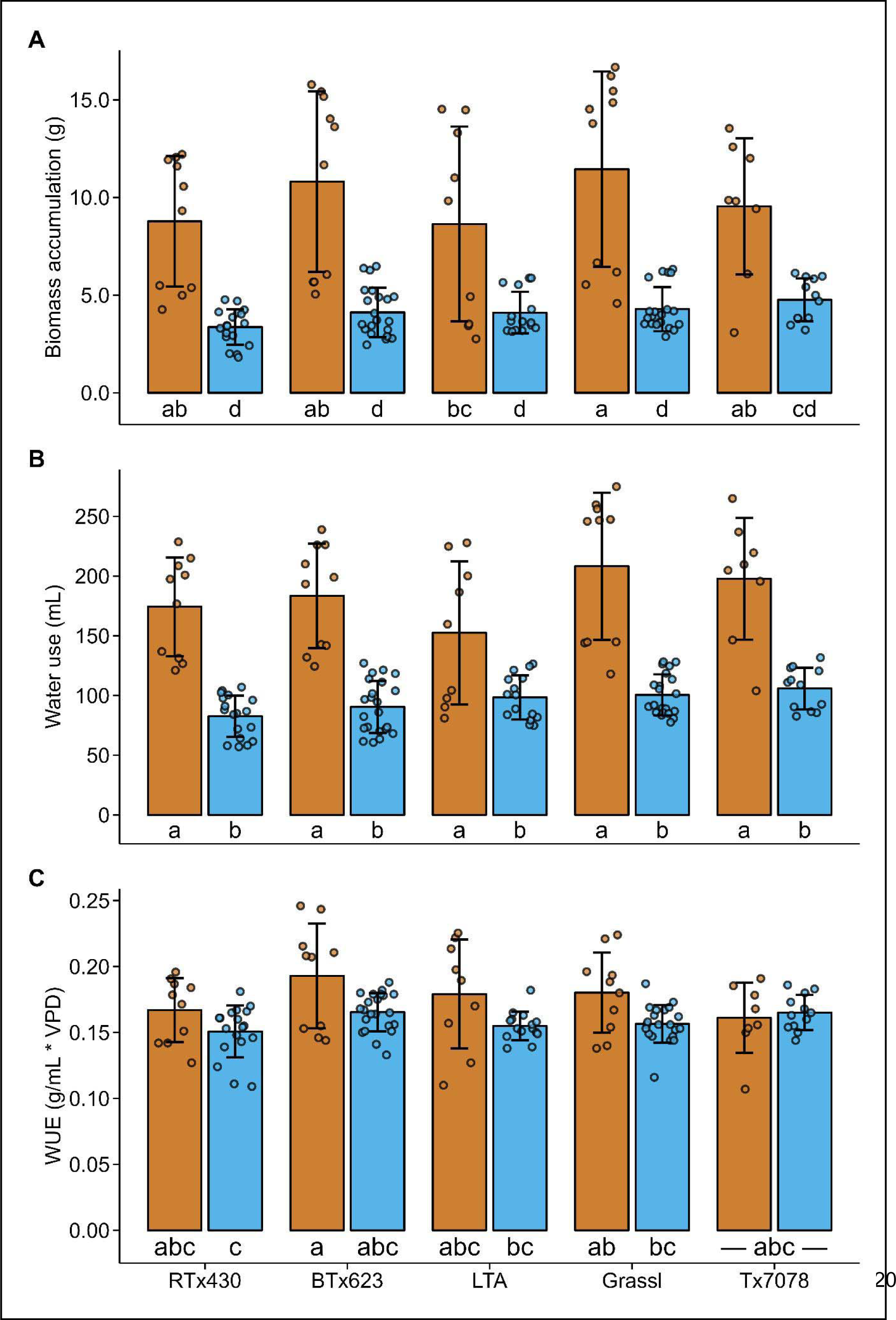
(A) Biomass accumulation, (B) water use, and (C) WUE of plants grown in soil (brown) or hydroponically (blue). Error bars represent the standard deviation (n = 8-22; N = 3-4). Letters represent significantly different groups as determined by two-way ANOVA followed by Tukey’s HSD test (alpha = 0.05). n, number of biological replicates per genotype and condition; N, number of independent experiments.

**Figure S3:**
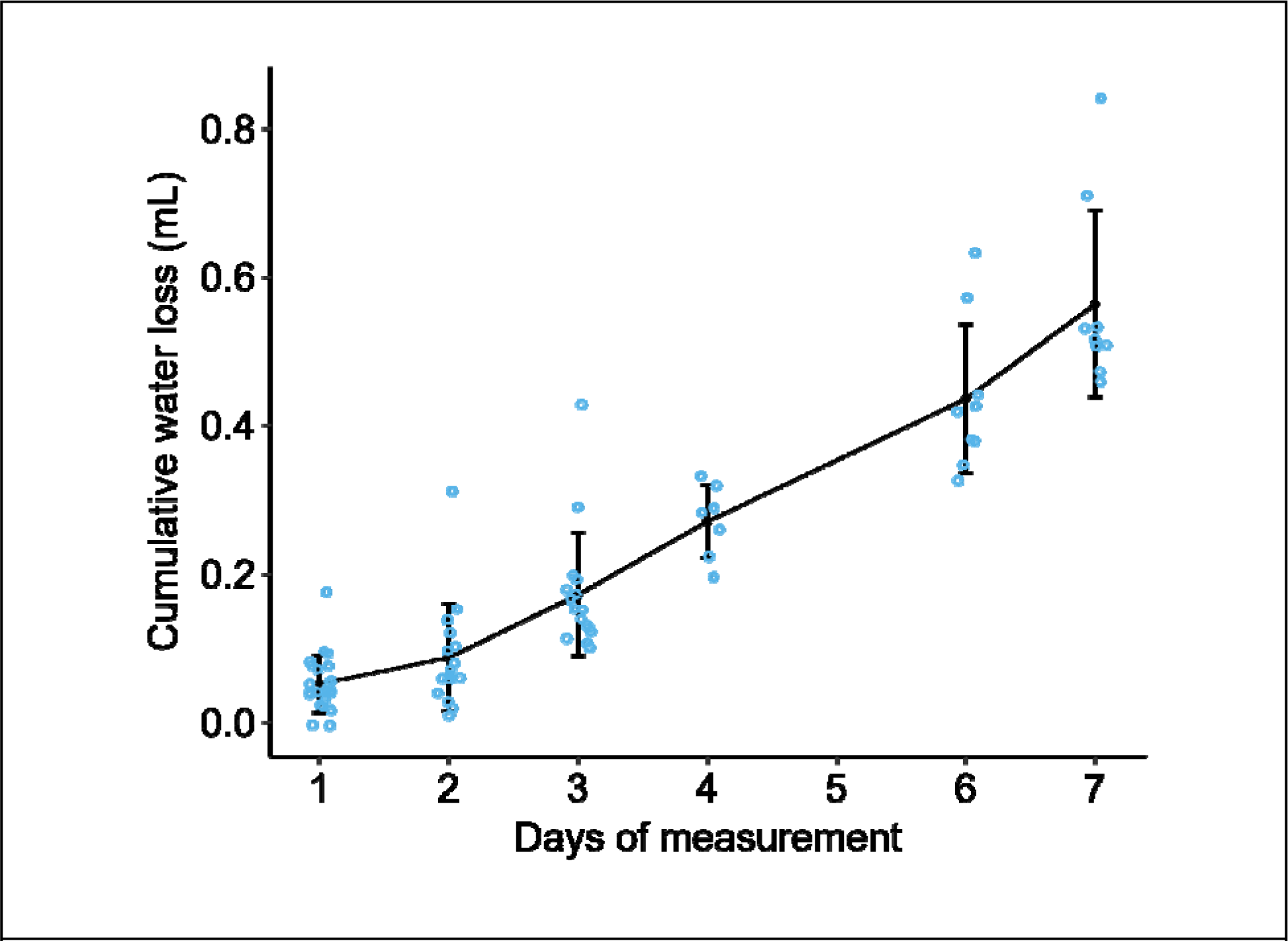
Water loss due to evaporation in closed hydroponic tubes. Black dots represent average cumulative evaporative water loss from closed tubes from which plant stems and roots were removed. Error bars represent the standard deviation (n = 7-23; N = 2-3). n, number of biological replicates per genotype and condition; N, number of independent experiments.

**Figure S4:**
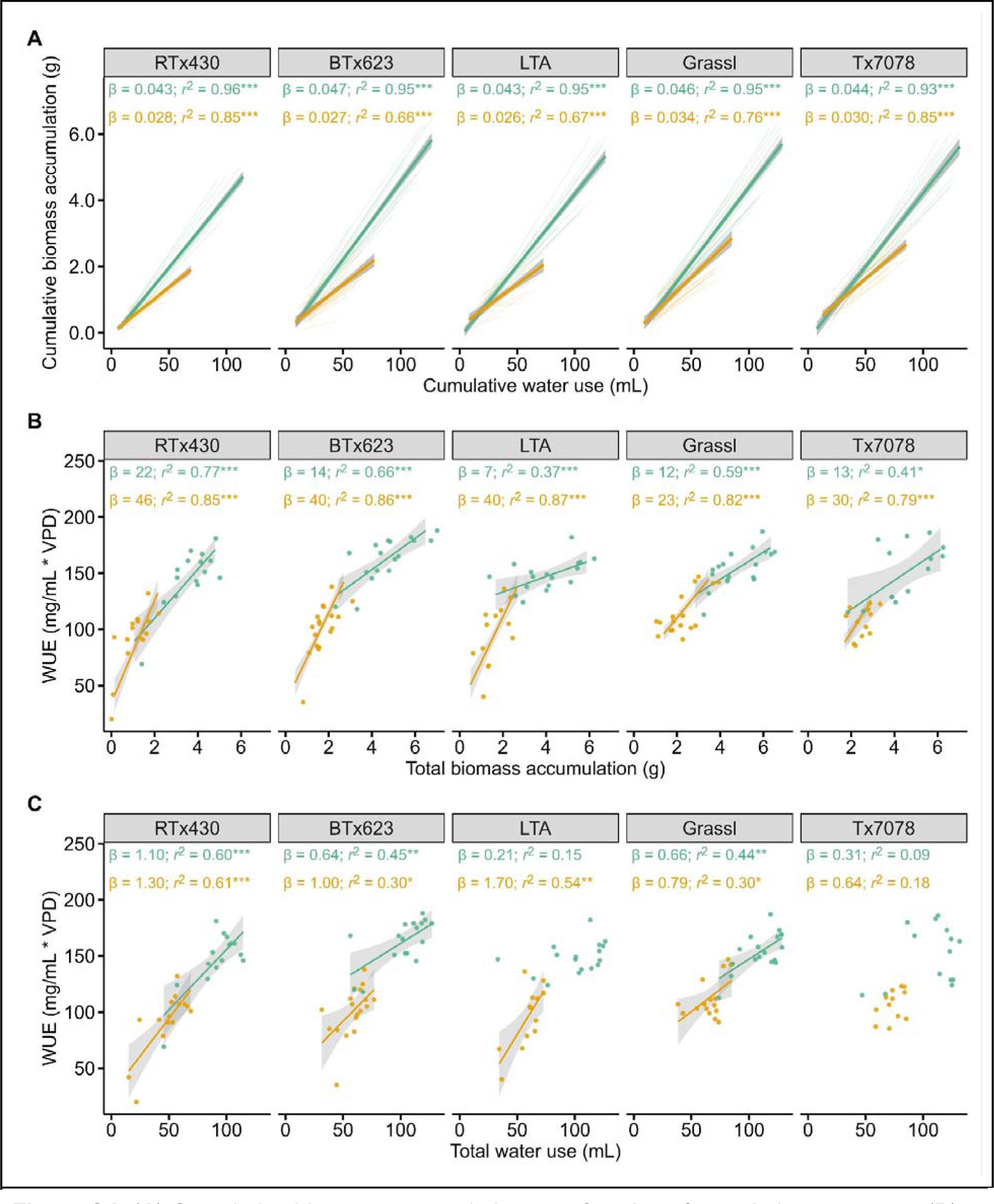
(**A**) Cumulative biomass accumulation as a function of cumulative water use, (**B**) WUE as a function of total (8-day) biomass accumulation, and **(C**) WUE as a function of total (8-day) water use in control (green) or mannitol-treated (orange) conditions (n = 12-18 per genotype, condition, and day; N = 4). Thick solid lines represent the treatment-level linear regression bounded by the 95% confidence interval. Thin lines in A represent linear regressions of individual samples. β values represent the slope (coefficient) of the linear regression. r^2^ values represent the square of the Pearson correlation coefficient of the linear regression. Asterisks indicate whether the linear relationship is statistically significant: * (p-value < 0.05), ** (p-value < 0.01), *** (p-value < 0.001). Absence of a regression line indicates a non-significant linear relationship (p-value > 0.05). n, number of biological replicates per genotype and condition; N, number of independent experiments.

